# Unravelling the diversity Behind the *Ophiocordyceps Unilateralis* Complex: Three new species of zombie-ant fungi from the Brazilian Amazon

**DOI:** 10.1101/003806

**Authors:** João P. M. Araújo, Harry C. Evans, David M. Geiser, William P. Mackay, David P. Hughes

## Abstract

In tropical forests, one of the most common relationships between parasites and insects is that between the fungus *Ophiocordyceps* (Ophiocordycipitaceae, Hypocreales, Ascomycota) and ants, especially within the tribe Camponotini. Here, we describe three new and host-specific species of the genus *Ophiocordyceps* on *Camponotus* ants from the central Amazonian region of Brazil, which can readily be separated using morphological traits, in particular, ascospore form and function. In addition, we use sequence data to infer phylogenetic relationships between these taxa and closely related species within the *Ophiocordyceps unilateralis* complex, as well as with other members of the family *Ophiocordycipitaceae*.

## Introduction

In tropical forests, social insects (ants, bees, termites and wasps) are the most abundant land-dwelling arthropods. Although they represent only 2% of the nearly 900,000 known insect species on Earth, they are estimated to compose more than half the biomass (Fittkau & Klinge 1973; Höldobler & Wilson 2009). One of the better known members within this group are the ants, which form a single family (Formicidae), with close to 13,000 species described (Agosti & Johnson 2009). Ants occupy a wide range of habitats from high canopy to the leaf litter, forming huge colonies comprising tens to hundreds of thousands to millions of individuals.

Ants are associated with and susceptible to a variety of parasites. Amongst these, one group is particularly well adapted to live in tropical forests and to exploit this ant abundance, entomopathogenic fungi of the genus *Ophiocordyceps* (Hypocreales; Ophiocordycipitaceae), currently comprising around 160 species (Robert et al. 2005; Sung et al. 2007). These parasites infect many different insects with a wide range of ecologies, from solitary wandering beetles to highly-organized ant societies. The orders infected are: Blattaria, Coleoptera, Dermaptera, Diptera, Hemiptera, Hymenoptera, Isoptera, Lepidoptera, Mantodea, Odonata and Orthoptera (Araújo & Hughes Preprint). The functional morphology of *Ophiocordyceps* is equally diverse and has been linked to the host’s ecology and biology (Evans et al. 2011).

Species of *Ophiocordyceps* were originally placed within *Cordyceps*, a genus established to accommodate fungal pathogens of arthropods bearing the sexual spore-producing structures on conspicuous stalks, arising from the host cadaver (Evans et al. 2011). However, due to the polyphyletic nature of *Cordyceps* – as evidenced by recent phylogenetic studies – species formerly assigned to the genus have now been reorganized into four genera (*Cordyceps, Elaphocordyceps, Metacordyceps* and *Ophiocordyceps)*, within three families (*Cordycipitaceae, Clavicipitaceae* and *Ophiocordycipitaceae)* (Sung et al. 2007).

Within the Formicidae, *Ophiocordyceps* infections have been reported from the basal primitive groups (Ponerines) through to modern genera, such as *Camponotus* (Evans & Samson, 1982; Sanjuan et al. 2001; Evans et al. 2011), and often occur as epizootics, killing large number of ants in small patches of forest (Andersen et al. 2009; Pontoppidan et al. 2009). Such events are pan-tropical with records from Asia, Australasia, Africa and the Americas (Evans 1974; Andrade 1980; Evans & Samson 1982, 1984; Evans 1988a, 2001; Kepler et al. 2011; Luangsa-ard et al. 2011; Kobmoo et al. 2012).

Entomopathogenic fungi infect their hosts following spore contact and subsequent germination on the cuticle. Typically, an adhesive pad (appressorium) is formed, whereby a germ tube penetrates the exoskeleton of the host. Upon reaching the haemocoel, the fungus proliferates as yeast-like cells (Evans 1988b). In the *O. unilateralis* complex, a series of synchronized events are triggered within the ant host in order to make it leave the colony, climb understorey shrubs to die in an elevated position - characteristically, biting the underside or edge of a leaf - and, once there, a spore-producing structure arises from the back of its head from which the ascospores are forcibly released at maturity (Andersen et al. 2009).

The taxonomy and evolutionary relationships of this important group of pathogens remain unclear. For many years it was suspected that *O. unilateralis*, originally described as *Torrubia unilateralis* (Tulasne & Tulasne 1865), represents a complex of species, based on macro-morphological variation within collections worldwide (Petch 1931; Kobayasi 1941; Mains 1958; Samson et al. 1982; Evans & Samson 1984). However, it was not until recently that species delimitation was proposed formally, when four new taxa were described from the Minas Gerais state of Brazil (Evans et al. 2011), and it was posited that each ant within the Camponotini would host a different species of *Ophiocordyceps.* Subsequent descriptions of new taxa, from both Thailand and Japan on both *Camponotus* and *Polyrhachis* hosts, are lending support to this hypothesis (Luangsa-ard et al. 2011; Kepler et al. 2011; Kobmoo et al. 2012) - and, even more recently, from the USA (de Bekker et al. 2014) - with the clear indication that there are still many species to be discovered.

During field surveys in the central Brazilian Amazon, we collected a range of *Camponotus* species killed by *Ophiocordyceps*, often in large numbers. Based on macro-morphological characters, all the specimens fell within *O. unilateralis sensu lato:* typically, comprising a stalk (stroma)arising from the dorsal neck (anterior of the pronotum); with an asexual morph (*Hirsutella*) occupying the terminal region and the sexual morph (ascoma) occurring as lateral cushions or plates. Here, we describe and characterize three new species, which can readily be separated using traditional micromorphology (Kobayasi 1941; Evans et al. 2011), as well as molecular data.

## Materials and Methods

### Sampling

Surveys were undertaken in the central Amazonian region of Brazil concentrating on four forest reserves (Figure 1): (A) Reserva Adolpho Ducke, ca. 10,000 ha (02°55’S, 59°59’W), adjacent to Manaus (Amazonas state) and composed of terra-firme forest, with plateaus, lowlands and campinarana vegetation, occurring on sandy soil across the Rio Negro basin; (B) Parque Nacional de Anavilhanas, an archipelago (2°23’S, 60°55’W) of more than 400 islands, 40 km up the Rio Negro from Manaus, colonized by Igapó forest (varzea), which is characterized by water-soaked soils with high acidity due to seasonal flooding; (C) Parque Nacional do Viruá, ca. 227,000 ha (01°42’ N, 61°10’ W), located near Caracaraí city (Roraima state), based on sandy soils with many lagoons and lowland Amazonian forest; (D) Estação Ecológica de Maracá, ca. 104,000 ha (3°22’N, 61°27’W), located 135 km from Boa Vista on an island of the Uraricoera River in Roraima state, containing both savanna and a mix of humid lowlands and plateaus (terra-firme).

Sampling protocol consisted of a careful inspection of soil, leaf litter, shrub leaves and tree trunks, up to 2m high. Infected ants, and the substrata they were attached to, were collected in plastic containers for transport to the laboratory and, wherever possible, examined the same day. During longer surveys, the samples were air-dried overnight to prevent mold growth. All specimens were photographed individually, using a Canon 60D camera fitted with a MP-E 65 mm (× 5) macro lens, equipped with a MT-24EX macro lite flash.

**Figure 1:**
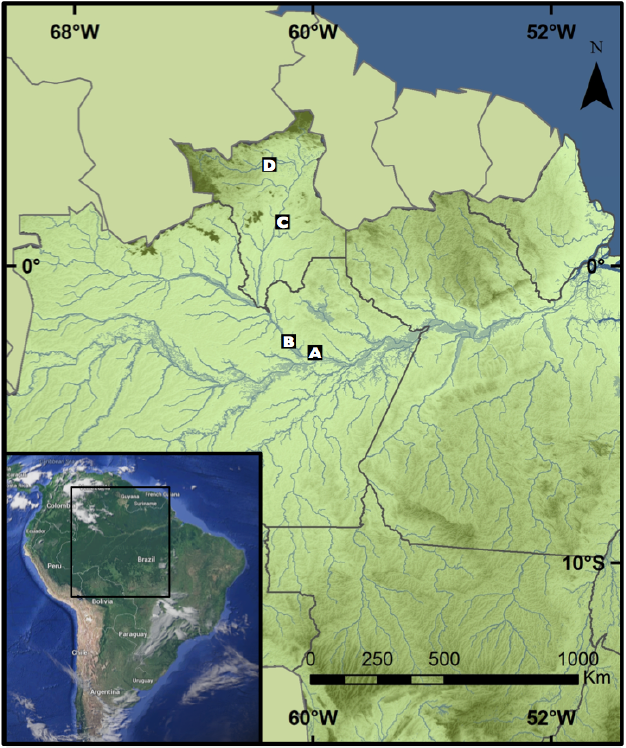
Map showing the forest reserves sampled in the central Brazilian Amazon: A) Reserva Adolpho Ducke; B) Parque Nacional de Anavilhanas; C) Parque Nacional do Viruá; D) Estação Ecológica de Maracá.

### Morphological Studies

Samples were screened using a stereoscopic microscope, and only mature fungal specimens were selected for further micro-morphological studies. In order to obtain ascospores, infected ants were attached to the lid of a plastic Petri plate (9-cm diam) using petroleum jelly, and suspended above a plate containing either distilled water agar (DWA) or potato-dextrose agar (PDA). Plates were transferred covered stands placed on the forest, subject to natural temperature and light fluctuations and examined daily for the presence of ascospores, which, after ejection from the ascomata, formed sub-hyaline halos on the agar surface. Freshly deposited ascospores were removed with a sterile hypodermic needle under a stereoscopic microscope, and mounted on a slide in lactofuchsin (0.1g of acid fuchsin in 100 ml of lactic acid) for light microscopy (Olympus BX61). A minimum of 50 naturally-released (mature) ascospores were measured for morphological comparison (Table 1). The remaining ascospores were left *in situ* on the agar surface and examined over a number of days in order to monitor germination events. For micro-morphology of the ascomata, either free-hand or cryo-sectioning (Leica CM1950 Cryostat) was used.

Permanent slides were deposited in the Entomological Collection at INPA (Instituto Nacional de Pesquisas da Amazônia, Manaus), with isotype collections held in Frost Entomological Museum at PSU (Penn State University). Permits for collecting and export were provided by SISBIO to JPMA (Authorizations 40496–1; 32435–1; 32410–1).

### DNA Extraction, PCR, Sequencing and Phylogenetic Analyses

DNA extractions were made using a small piece of the ascomata and hyphae within the ant’s abdomen. The tissue was placed with two metal balls (1/8″) in a 2 ml Eppendorf tube, frozen in liquid nitrogen and broken into powder using Tissuelyzer (Qiagen) for 1 min. Afterwards, with the fungal tissue still frozen, 0.8 ml of DNA extraction buffer at 50°C (1% SDS, 0.024 g/ml PAS, 0.2 ml/ml RNB, distilled water) and 0.8 ml phenol-chloroform were added, incubated at 60°C for 10 min and centrifuged for 10 min at 10,000 rpm. The water phase was transferred to a new 2 ml tube (about 0.8 ml), 0.5 ml of chloroform was added and the contents mixed by inverting, and then centrifuged for 10 min at 10,000 rpm. The supernatant was transferred to a new tube, 80% of the volume of isopropanol was added and both mixed by inverting, and then centrifuged for 10 min at max speed (14,000 rpm). The supernatant was removed and the pellet formed at the bottom was washed with 0.5 ml 70% ethanol 200 proof pure ethanol (Koptec), centrifuged for 5 min at 14,000 rpm, and the supernatant. The tube with the pellet was air-dried and dissolved in 30 μl TE buffer.

Approximately 1,030 bp of nu-SSU, 770 bp of nu-LSU and 550 bp of ITS were amplified by PCR. The nu-SSU amplification was performed using two overlapping sets of primers, NS1/ SR7 (R. Vilgalys, 5’-GTTCAACTACGAGCTTTTTAA-3’) and NS4/ NS3 (White et al. 1990). The nu-LSU amplification was amplified with the primers LR5/LR0R (Vilgalys and Sun, 1994) and for ITS amplification, ITS1F (Gardes & Bruns 1993) and ITS4 (White et al. 1990) were used. PCR was performed in a T-3000 Thermocycler (Biometra GmbH, Göttingen, Germany) with the following mix: 2.5 μl PCR buffer, 0.5 μl dNTP, 1.5 MgCl, 0.5 each primer, 1 μl template, O.1 Invitrogen Taq Platinum Polymerase and 18.4 μl Gibco UltraPure Distilled Water. PCR products were cleaned using the Qiagen PCR Purification Kit following the manufacturer’s instructions and sequenced using the Genomics Core Facility service at PSU.

Sequences were edited manually using Sequencher version 4.7 (Gene Codes Corp., Ann Arbor, MI, USA). Sequences generated for this study were compared with closely related species sequences available on GenBank. Herbarium numbers and GenBank accession numbers are provided in Table 1. The alignment was obtained by MUSCLE algorithms using Mega 5 (Kumar et al. 2012) and gaps were treated as missing data. Maximum Likelihood and Maximum Parsimony analyses were conducted using Mega 5 (Kumar et al. 2012) with 1,000 bootstrap replicates on a concatenated dataset containing all three genes.

## Results

### Taxonomy

***Ophiocordyceps camponoti-atricipis*** Araújo, H.C. Evans & D.P. Hughes sp. nov.

#### IF 550743

Etymology: Named after the ant host, *Camponotus* (*Myrmothrix*) *atriceps* Mayr.

Type: Brazil. Amazonas: Reserva Adolpho Ducke, 100 m, 10 Jan 2012 J.P.M. Araújo & H.C. Evans, A-35, on *Camponotus atriceps* Smith (holotype INPA #; isotype INPA #, FROST #).

External mycelium abundant, covering most of the host, produced from all orifices and sutures; initially white, turning light-brown. Stroma single, produced from dorsal pronotum, averaging 15-20 mm, up to 25 mm in length, cylindrical, velvety and ginger brown at the base, becoming cream-pinkish towards apex; fertile region of lateral cushions, 1-2, hemispherical, chocolate brown, darkening with age, variable in size, averaging 1.5 × 0.5−0.8 mm. Perithecia immersed to partially erumpent, flask-shaped, 240−280 × 100−150 μm, with short, exposed neck or ostiole. Asci 8-spored, hyaline, cylindrical to clavate, 110−140 × (4.5−) 6−6.5 (−8) μm; prominent cap, (3.5−) 5 × 5.5 (−6.5) μm. Ascospores hyaline, thin-walled, vermiform (75−) 80−85 (−100) × (2−) 3 (−3.5) μm, 5− septate, sinuous to curved, never straight at maturity, rounded to acute apex.

##### Asexual Morph

*Hirsutella*-A type only, produced on the upper stromatal surface; phialides cylindrical to lageniform, 5−7 × 2−3 μm, tapering to a long neck, 5−11 μm; conidia not seen. This asexual morph occurs in all the species included here and is not considered to be critical for species separation. Hence, it is not analyzed in detail.

##### Germination Process

The released ascospores germinated within 24-48 h, producing 1-2, uniformly straight, thread-like structures (capillicondiophores); typically of uniform length and averaging 55 μm; bearing a single terminal spore (capilliconidium), hyaline, smooth-walled, allantoid and tapering at the ends at maturity, 10-11 × 2-2.5 μm, narrowing apically.

##### Habitat

Hosts commonly found biting the apical part of palm-fronds, but also on dicot leaves, rarely on palm-spines; abundant mycelial growth from the mouthparts helping to fix the host to the leaf, in addition to the locked jaws.

**Figure 2:**
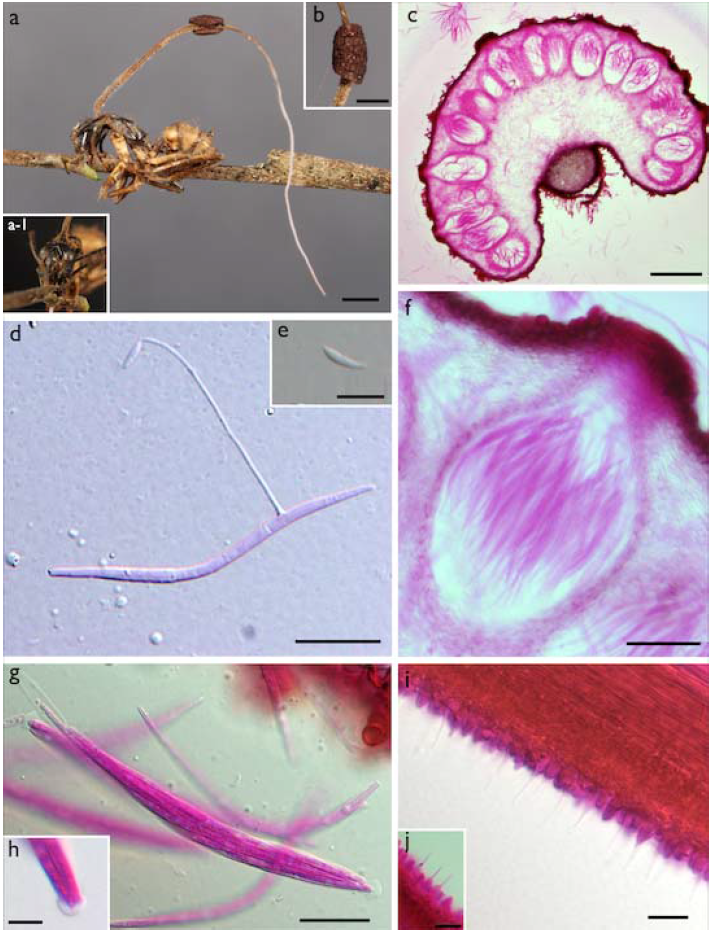
*Ophiocordyceps camponoti-atricipis* **a)** Single stroma, characteristic of *Ophiocordyceps unilateralis sensu lato*, with a single lateral ascoma, arising anterior to pronotum of *Camponotus atriceps*, firmly attached to apex of palm frond (bar = 1.5 mm); **a-1**) Detail of biting ant ; **b)** Detail of fertile region (ascoma) (bar = 0.8 mm); **c)** Section through ascoma showing the mainly immersed perithecial arrangement (bar = 200 μm); **d)** Ascospore with a needle-like outgrowth (capilliconidiophore) producing terminal conidium (bar = 20 μm); **e)** Close-up of conidium (bar = 10 μm); **f)** Close-up of perithecium (bar = 50 μm); **g)** Ascus, clavate in shape and with a prominent cap (bar = 20 μm); **h)** Detail of the ascus cap (bar = 5 μm); **i)** Section of upper part of stroma showing asexual morph (*Hirsutella*-A type), with a palisade of subulate phialides (bar = 10 μm); **j)** Close up (bar = 10 μm).

***Ophiocordyceps camponoti-bispinosi*** Araújo, H.C. Evans & D.P. Hughes sp. nov.

#### IF 550744

Etymology: Named after the ant host, *Camponotus (Myrmocladoecus) bispinosus* Mayr.

Type: Brazil. Amazonas: Reserva Adolpho Ducke, 100 m, 15 Jan 2012, J.P.M. Araújo & H. C. Evans, B-66, on *Camponotus bispinosus* Mayr (holotype: INPA #; isotype FROST #).

Mycelium dark-brown to black; forming aggregations of hyphae on the intersegmental membranes. Stroma single, produced anterior to pronotum, averaging 5-7 mm in length; cylindrical, black, with a cream-white swollen terminal part; fertile ascomatal region lateral, forming a single globose cushion, initially dark brown becoming black with age, averaging 0.8 × 0.4−0.7 mm. Perithecia immersed to slightly erumpent, globose to flask-shaped, 250-290 × 150−170 μm, with a short neck. Asci 8-spored, hyaline, cylindrical to clavate, (90−) 110−130 × (7−) 8−8.5 (−9.5) μm; with a small but prominent cap, 3.5 × 4.5 μm. Ascospores hyaline, thin-walled, cylindrical, (60−) 70−75 (80) × 4.5−5 (−6), 4−5 septate, round and slightly narrow at the apices.

##### Asexual Morph

*Hirsutella*-A type only, produced at the swollen upper part of stroma; phialides cylindrical to lageniform, 6 × 2.5−3 μm, tapering to a long hair-like neck, 10−16 μm length; conidia narrow limoniform, averaging 6−7 × 2 μm, with a distinct tail.

##### Germination Process

Spores germinated 48-72 h after release, consistently producing a single, straight, robust capilliconidiophore, (50−) 65 (−80) μm, bearing a single terminal capilliconidium, 10−11 × 3−4 μm, hyaline, smooth-walled, slightly truncate at the base.

##### Habitat

Often biting palm-tree parts, commonly on spines or tip-edge of the frond; mycelium growing from mouth, attaching the ant to the substrate; rarely on broad-leaved plants.

**Figure 3:**
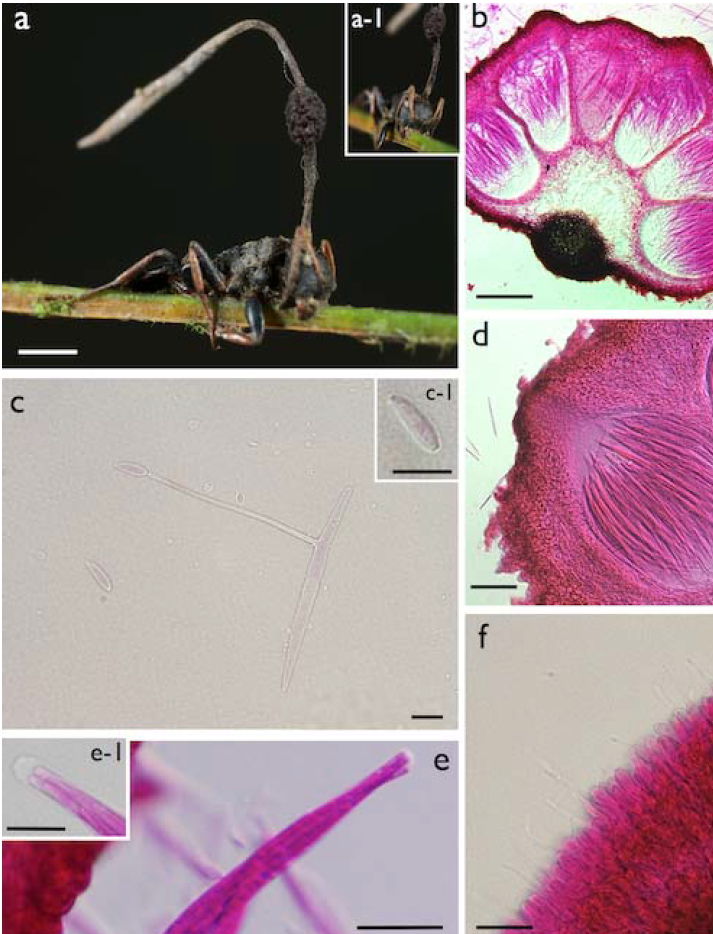
***Ophiocordyceps camponoti-bispinosi* a)** Infected *Camponotus bispinosus* biting into tip of palm leaf **a-1**) Detail showing biting behavior and ascoma; **b)** Section through ascoma showing perithecial arrangement (bar = 100 μm); c) Ascospore after 48 h germinating with a single capilliconidiophore with a capilliconidium at the tip (bar = 10 μm); c−1) Detail of the capilliconidium (bar = 10 μm); d) Section of perithecium showing the arrangement of asci (bar = 50 μm); e) Ascus showing the ascospore arrangement (bar = 20 μm); e−1) Detail showing the prominent ascus cap (bar = 10 μm); f) Section of the swollen stromatal tip showing the asexual morph (*Hirsutella*-A)(bar = 10 μm).

***Ophiocordyceps camponoti-indiani*** Araújo, H.C. Evans & D.P. Hughes sp. nov.

#### IF 550745

Etymology: Named after the ant host, *Camponotus (Tanaemyrmex) indianus* Forel.

Type: Brazil. Roraima: Parque Nacional do Viruá, 9 Feb 2012, V-15, on *Camponotus indianus.*

Mycelium aggregations arising from all inter-segmental membranes (sutures), ginger in color. Stromata multiple, arising from dorsal, right and left sides of pronotum, and leg joints, (1.5−) 8−10 (16) × 0.3−0.4 (−1) mm, ginger at the base becoming purplish-cream towards the apex. Ascomata produced only on the pronotal stromata, never from those on legs; lateral cushions, 1−4, hemispherical, chocolate to dark brown with age. Perithecia immersed to semi-erumpent, ovoid to flask-shaped, 230−310 × 120−175 μm, with a short, exposed neck. Asci 8-spored, hyaline, cylindrical (135−) 170 (−190) × 8.5 (11.5) μm; cap prominent, 4.5 × 5 μm; Ascospores hyaline, thin-walled, cylindrical, (60−) 75 (−80) × (3.5−) 4.5 (−5), 5-septate, occasionally swollen, narrowing to acute tips at both ends.

##### Asexual Morph

*Hirsutella*-A type associated with the apical region of all stromata; phialides cylindrical to lageniform, 7.5 (−9.5) × 3.5 μm, tapering to a long hair-like neck, 6.5−11 μm in length, conidia not seen. *Hirsutella-C* type (= *H. sporodochialis-* type) produced from ginger cushions (sporodochia) on legs and antennal joints: phialides hyaline and subulate at the base, robust; no conidia observed.

##### Germination Process

Ascospores germinated 48 h after release; typically, producing 2, rarely 3, hair-like capilliconidiophores, 120−130 μm in length; bearing a single terminal conidium, 13–14 μm biguttulate, fusoid, narrowing apically.

**Figure 4:**
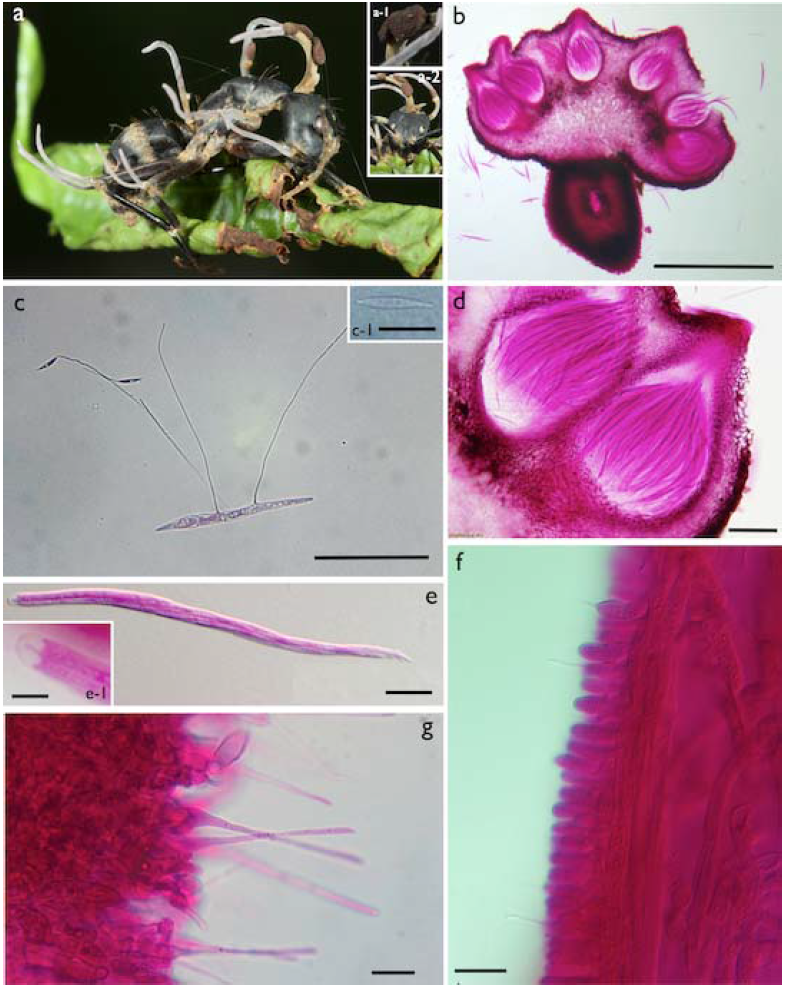
***Ophiocordyceps camponoti-indiani* a)** *Camponotus indianus* biting into a leaf, several stromata arising from dorsal pronotum, mesonotum and leg joints, with a characteristic purplish coloration. **a-1)** lateral, fertile cushion (ascoma) ; **a-2)** Close up of the dead ant’s head showing the biting behavior. **b)** Section through ascoma showing perithecial arrangement (bar = 500 μm); **c)** Ascospore after 24 h, with very long capilliconidiophores (1-3) with capilliconidia at the tip (bar = 50 μm); **c-1)** Detail of fusoid capilliconidium (bar = 10 μm); **d)** Close up of perithecia showing asci arrangement and the semi-erumpent ostiole (bar = 50 μm); **e)** Ascus showing the spiral arrangement of ascospores (bar = 20 μm); e−1) Ascus cap detail (bar = 5 μm); **f)** Section of upper part of stroma showing asexual morph (*Hirsutella-A* type), with long-necked phialides (bar = 10 μm); **g)** Phialides formed as mycelial cushions (sporodochia) on leg joints and antenna (*Hirsutella* C-type) (bar = 10 μm).

**Table 1.** Comparison of main morphological characters of the new *Ophiocordyceps* species and closely related species.

|  | Host | Stroma (mm) | Capilliconidiophore length (µm) | Ascospore dimension (µm) | Hirsutella Type |  |  |
| --- | --- | --- | --- | --- | --- | --- | --- |
|  |  |  |  |  | A | B | C |
| *O. camponoti-atricipis* * | *Camponotus atriceps* | 15-20 (-25) | 55 | 80-85 x 3<br>**(5) | + | - | - |
| *O. camponoti-bispinosi* * | *C. bispinosus* | 5-7 | 65 | 70-75 x 4.5<br>(4-5) | + | - | - |
| *O. camponoti-indiani* * | *C. indianus* | 8-10 (-16) | 130 | 75 x 4.5<br>(5) | + | - | + |
| *O. camponoti-rufipedis* | *C. rufipes* | 5-8 (-15) | 60-70 (-80) | 80-95 - 2-3<br>(4-7) | + | - | - |
| *O. camponoti-balzani* | *C. balzani* | similar to *O.c-rufipedis* | - | 135-175 x 4-5 (14-22) | + | - | + |
| *O. camponoti-melanotici* | *C. melanoticus* | similar to *O.c-rufipedis* | - | 170-210 x 4-5 (27-35) | + | - | - |
| *O. camponoti-novogranadensis* | *C. novogranadensis* | similar to *O.c-rufipedis* | 20-25 | 75-95 x 2.5-3.5 (5-10) | + | + | - |
| *O. halabalaensis* | *C. gigas* | 6.5-18 | - | 60-75 x 3-5<br>("multi-septate") | + | - | - |

\*New species
\*\*Septation

**Table 2.** Taxon, specimen voucher, host affiliation and sequence information for specimens used in this study

| Taxon | Specimen voucher | Host | GenBank accession numbers |  |  |
| --- | --- | --- | --- | --- | --- |
|  |  |  | nu-SSU | nu-LSU | ITS |
| *Ophiocordyceps camponoti-atricipis* | Will be provided by INPA | *Camponotus atriceps* (Ant-Hymenoptera) | KJ953886 | KJ953889 | KJ953892 |
| *O. camponoti-bispinosi* | Will be provided by INPA | *Camponotus bispinosus* (Ant-Hymenoptera) | KJ953887 | KJ953890 | KJ953893 |
| *O. camponoti-indiani* | Will be provided by INPA | *Camponotus indianus* (Ant-Hymenoptera) | KJ953888 | KJ953891 | KJ953894 |
| *O. irangiensis* | OSC 128577 | Ant (Hymenoptera) | DQ522546 | DQ518760 | JN049823 |
| *O. kniphofioides* | Ophkni790 | *Cephalotes atratus* (Ant - Hymenoptera) | KC610789 | KC610767 |  |
| *O. nigrella* | EFCC 9247 | Scarabaeidae larva (Coleoptera) | EF468963 | EF468818 | JN049853 |
| *O. nutans* | OSC 110994 | Pentatomidae (Hemiptera) | DQ522549 | DQ518763 |  |
| *O. pulvinata* | TNS-F 30044 | *Camponotus obscuripes* (Ant - Hymenoptera) | GU904208 |  | AB721302 |
| *O. sphecocephala* | OSC 110998 | Wasp (Hymenoptera) | DQ522551 | DQ518765 |  |
| *O. stylophora* | OSC 111000 | Elaterid larva (Coleoptera) | DQ522552 | DQ518766 |  |
| *O. unilateralis* | OSC 128574 | Ant (*Camponotus* sp.) | DQ522554 | DQ518768 |  |
| *Tolypocladium capitatum* | NBRC 106327 | Fungi (*Elaphomyces* sp.) | JN941737 | JN941404 | JN943317 |

## Discussion

### Morphology

The three new species proposed herein differ considerably from other *Ophiocordyceps* species described previously. *Ophiocordyceps camponoti-atricipis, O. camponoti-bispinosi* and *O. camponoti-indiani* share common traits, such as *Camponotus* ants as the host, a stroma arising from the intersegmental region anterior to the pronotum, bearing one or more brown to blackish cushions attached laterally. Thus, they could readily be grouped within the *O. unilateralis s.l.* complex of species, based purely on macro-morphology. Further micro-morphological analyses in the lab revealed clear differences between our collections and previous species belonging to the *O. unilateralis* species complex. The ascospores - including their germination and growth behavior - and the host association were the most evident characters to distinguish the three new species. In addition, other features - such as ascoma, perithecia, asci, anamorphic morphology - were also examined and used for comparison and further separation from closely related species within the *O. unilateralis* complex (Figures 1–3; Table 1).

Using macro-morphological characters, *O. camponoti-indiani* is the only species that exhibits multiple (from 3 up to 12) pale-purplish stromata arising from several intersegmental membranes on legs and thorax. The stromatal surface is covered by *Hirsutella* phialides, producing conidia up to the tip, especially on sexually immature specimens (Fig. 4-f). A similar species has been described from Thailand, *O. halabalaensis* (Luangsa-ard et al. 2011), which likewise exhibits multiple stromata, but never more than three. However, no ascospore behavior data is available for *O. halabalaensis* to be compared to *O. camponoti-indiani. Ophiocordyceps camponoti-atricipis* and *O. bispinosi* exhibit the classic “unilateralis” macro-morphology, which is a single stroma arising between the head and dorsal pronotum bearing a laterally attached ascomatal cushion. *O. camponoti-atricipis* produces a long and slender stroma, which is chocolate brown and velvety from the bottom up to the ascoma insertion, becoming pale-pinkish and glabrous above and gradually forming a *Hirsutella*-A layer, producing conidia up to the tip (Fig. 2-j). *O. bispinosi* produces a much shorter black stroma, often swelling and becoming pale-cream in the last third (Figure 3-a), with a *Hirsutella-A* layer producing conidia (Fig. 3-h).

Microscopic features are also markedly different between the species. As with other taxa in the *O. unilateralis* clade proposed by Sung et al. (2007), the ascospores of all the new species do not disarticulate into part-spores, but they can readily be delimited on shape, size and septation (Table 1). In addition, their germination behavior can also be used to separate them, as reported for other species within the complex (Evans et al. 2011). *O. camponoti-indiani* ascospores germinate with one to three long capilliconidiophores that reach up to 130 μm (Figure 4-c), contrasting with the fewer and shorter capillioconidiophores in *O. camponoti-atricipis* (producing 1-2, up to 55 μm long) and *O. camponoti-bispinosi* (single and robust, averaging 65 μm in length) (Figures 2-d; 3-d). In addition, the shapes of the capilliconidia are also distinct between the three species (Figures. 2-e; 3-c1; 4-c1). Capilliconidiophore variation among the species, especially the length, may reflect an adaptation to the host biology and further studies aim to address this question. Our results confirm that it is justifiable to split *O. unilateralis s.l.* into many taxa based on host specificity, potentially, unraveling a huge diversity of undocumented species.

### Phylogenetic Relationships

Besides the morphological examination, we also investigated the new species at the molecular level. We used a dataset of three genes (nu-SSU, nu-LSU and ITS1-5.8S- ITS2) to construct a tree with *O. camponoti-atricipis, O. camponoti-bispinosi, O. camponoti-indiani*, as well as previously studied species within Ophiocordycipitaceae, including members of the unilateralis complex (O. *unilateralis s.s.* and *O. pulvinata).* Our findings corroborate the morphological and ecological data that the species proposed here belong to *O. unilateralis s.l.* and comprise three novel species of *Ophiocordyceps* on *Camponotus* ants (Figure 5). Further studies, including more genes and more samples, are necessary to elucidate the relationship between the *Ophiocordyceps* species pathogenic on ants and other Hypocreales species.

**Figure 5:**
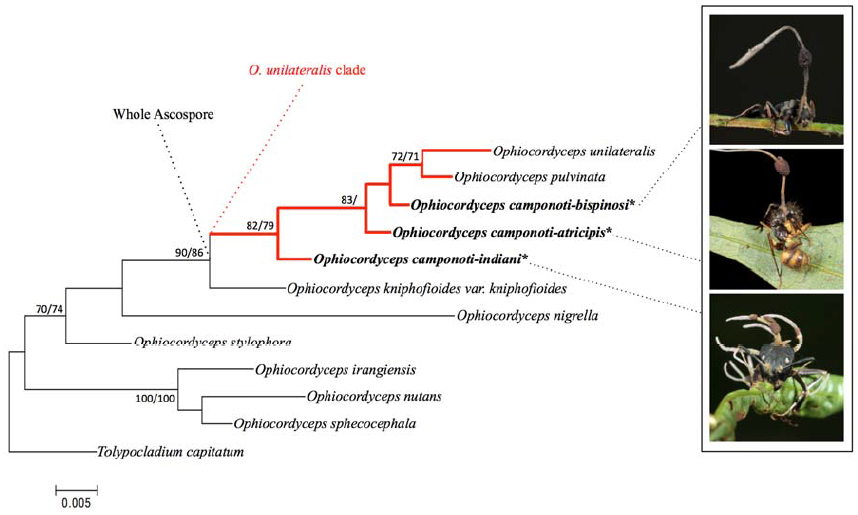
Maximum-likelihood tree obtained from a concatenated dataset of three genes (nu-SSU, nu-LSU, ITS) showing the placement of *O.camponoti-atricipis, O. camponoti-bispinosi* and *O. camponoti-indiani* within *Ophiocordyceps unilateralis* complex and relative to other Ophiocordycipitaceae species. Numbers above branches indicate bootstrap scores >70 (ML/MP).

### Ecological Aspects

The species described in the present study are currently known only from the Brazilian Amazon. Infected dead ants were collected biting onto shrub leaves and palm fronds, especially near the tips, as well as on palm spines and epiphytes, and, occasionally, on slender climbing stems or lianas. We observed that early in the morning, droplets of dew are formed on the leaf tips (especially of palm leaves), perhaps providing a daily supply of water for fungal development, even in the extended dry season. Thus, the adaptation of dying in this specific location may be advantageous for continual and consistent fungal development during the year. The same death position and substrate was also reported for *O. halabalaensis* in Thailand (Luangsa-ard *et al.* 2011).

*O. camponoti-atricipis* and *O. camponoti-bispinosi* were commonly found at multiple sites in all the four areas visited during this study, as well as in other areas previously visited across the Brazilian Amazon. The clusters of infections were composed of either one or multiple species of *O. unilateralis s.l.* in the same area, varying from just a few (i.e. 3-5) up to dozens of individuals. In some areas, the density was so high that it was possible to find ants biting into the same leaf or even into other infected ants (Figure 8). In contrast, the 19 *O. camponoti-indiani* specimens were all collected at the Parque Nacional do Viruá on a single occasion from one small site (~10 m^2^).

**Figure 7:**
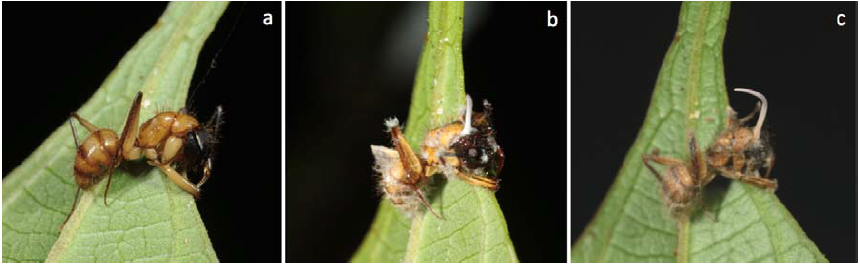
Infected *Ophiocordyceps camponoti-atricipis* showing initial development. **a)** Day 1 (24^th^ March 2011): Ant attaching to the leaf and dying a few hours later; **b)** Day 3: Cottony white fungal mycelium arises from ant sutures and joints, the stroma (synnema) emerges from behind the ant head; **c)** Day 5: the covering mycelium becomes light brown and the pink-tipped stroma continues to grow. In 2-3 weeks, the ascoma forms and matures over time depending on climatic conditions.

**Figure 8:**
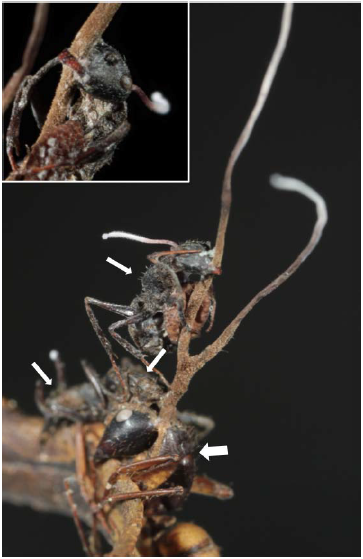
Unusual aggregation of different ant species biting on the same leaf and even onto the stoma from another infected ant. Arrows show the four different ants (two species) dead at the same spot.

**Figure 9:**
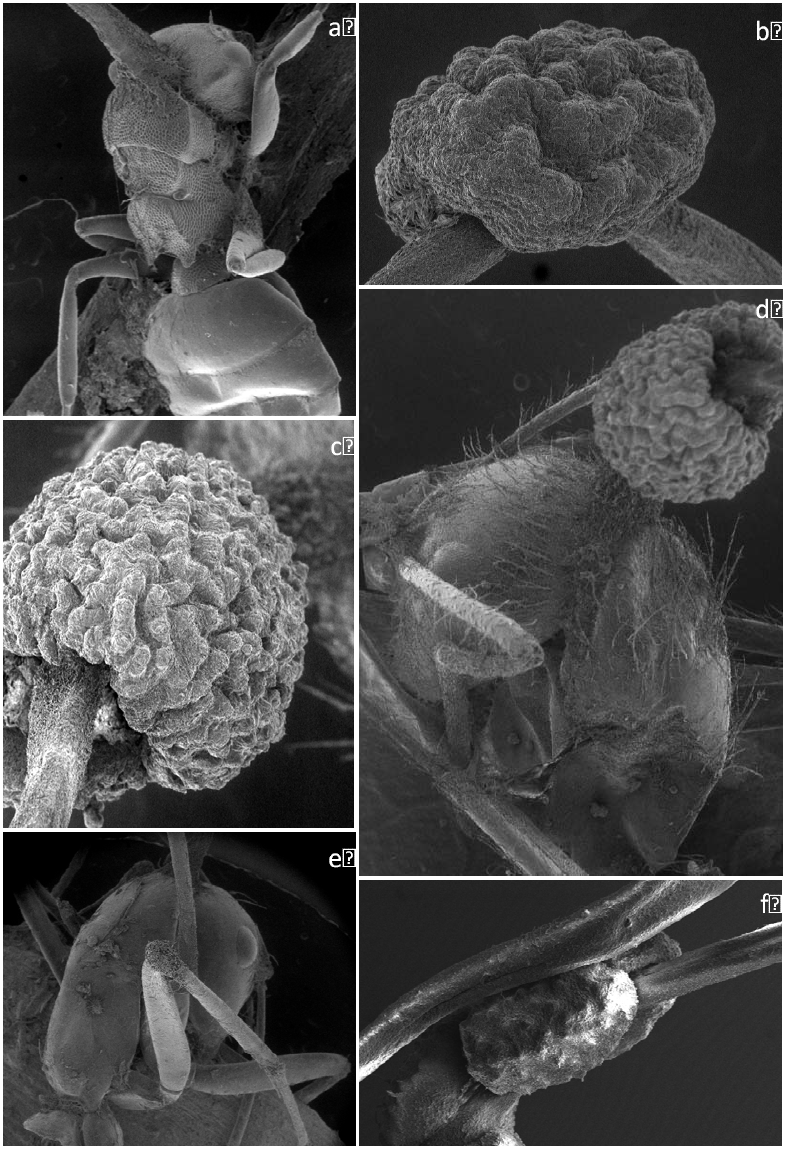
Scanning Electron Micrographs (SEM) of the infected ants. a) *C. bispinosus* infected by *O. camponoti-bispinosi;* b) Close-up of the *O. camponoti-bispori* ascoma; c) close up of the *O. camponoti-atricipis* ascoma; d) infected *C. atriceps*; e) infected *C. indianus* f) close up of *O. camponoti-indiani* ascoma.

## Conclusions

Ants of the genus *Camponotus* are parasitized by specialist fungal pathogens of the genus *Ophiocordyceps*, and this parasite-host association is especially common in tropical and sub-tropical forests. Based on macro-morphology, these pathogens – or, zombie-ant fungi - can all be assigned to *O. unilateralis s.l.* However, it has been posited that each species of *Camponotus* is parasitized by a host-specific species of the fungus and that these taxa can be separated on micro-morphology (Evans et al. 2011). Here, based on collections from central Amazonia, we show that this hypothesis is robust, particularly when the form and function of mature and living ascospores are compared. This has also been supported by molecular evidence and, therefore, we are confident that many more (perhaps, hundreds) new *Camponotus-Ophiocordyceps* associations remain to be discovered worldwide.

## Acknowledgements

We are grateful to Fabrício Baccaro (UFAM), Ricardo Braga-Neto (INPA) and Carlos Nogueira (INPA) for assistance in the field. Our special thanks to Márcio Oliveira and Thiago Mahlmann for the assistance with the permissions at INPA Entomological Collection…….

